# Osteoprotegerin is elevated in pulmonary fibrosis and associates with IPF progression

**DOI:** 10.1101/2020.12.02.408062

**Authors:** H. Habibie, Kurnia S.S. Putri, Carian E. Boorsma, David M. Brass, Peter Heukels, Marlies Wijsenbeek, Mirjam Kool, Maarten van den Berge, Theo Borghuis, Corry-Anke Brandsma, C Tji Gan, Peter Olinga, Wim Timens, Nicolas Kahn, Michael Kreuter, Janette K. Burgess, Barbro N. Melgert

**Author notes:** Correspondence should be addressed to: Prof. Dr. Barbro N. Melgert., Department of Molecular Pharmacology, Antonius Deusinglaan 1, 9713 AV Groningen, The Netherlands.

## Abstract

Osteoprotegerin (OPG), a decoy receptor for receptor activator of NF-kB ligand (RANKL), is used as a biomarker for assessing severity of liver fibrosis. However, its expression and role in pulmonary fibrosis are unknown. We hypothesized that OPG also has a role in pulmonary fibrosis.

Human and mouse control and fibrotic lung tissue were used to examine OPG expression, and mouse precision-cut lung slices to study OPG regulation in pulmonary fibrosis. Serum from idiopathic pulmonary fibrosis (IPF) patients and controls was analysed to investigate whether OPG levels correlate with disease status as measured by lung function.

OPG-protein levels were significantly higher in mouse and human fibrotic lung tissue compared to control. OPG-mRNA and protein production were induced in mouse precision-cut-lung slices upon TGFβ stimulation and could be inhibited with galunisertib, a TGFβ receptor kinase inhibitor. OPG-protein levels in fibrotic mouse lung tissue correlated with degree of fibrosis. Isolated lung fibroblasts from IPF patients had higher OPG-protein levels than control fibroblasts. Serum OPG levels in IPF patients, at first presentation, negatively correlated with diffusing capacity to carbon monoxide. Finally, serum OPG levels higher than 1234 pg/ml at first presentation were associated with progression of disease in IPF patients.

In conclusion, OPG is produced in lung tissue, associates with fibrosis, and may be a potential prognostic biomarker for IPF disease progression. Validation in a larger cohort is warranted to further explore the role of OPG in pulmonary fibrosis and its potential for assessing the prognosis of fibrotic lung disease in individual patients.

**Take home message:** Osteoprotegerin is present in fibrotic lung tissue and high serum levels correlate with low lung function and IPF disease progression in this small study, indicating osteoprotegerin may have value as a biomarker to predict IPF progression

## INTRODUCTION

Pulmonary fibrosis is characterized by extracellular matrix (ECM) deposition and progressive destruction of lung architecture that leads to impaired lung function with high mortality rates [1-3]. Idiopathic pulmonary fibrosis (IPF) is the most severe form of pulmonary fibrosis and has the highest mortality [4, 5]. One of the challenges in identifying and managing IPF is the lack of biomarkers to assist diagnosing the disease and predicting progression and therefore most patients experience a delayed diagnosis [6]. Having reliable biomarkers to diagnose IPF faster and to track disease progression could improve management of this disease and identification of optimal timing for referral for lung transplantation [7].

Osteoprotegerin (OPG), a decoy receptor for receptor activator of nuclear factor κB ligand (RANKL) and tumour necrosis factor-related apoptosis-inducing ligand (TRAIL) [8], is prominently expressed in bone tissue and best known for its role in the regulation of bone matrix [9, 10]. We recently showed high OPG expression in liver fibrosis with profibrotic actions of this molecule and the possible use of OPG in assessing treatment responses to antifibrotic drugs [11]. Others have shown associations with fibrotic processes in heart [12] and vasculature [13] and clinically OPG is used in a panel of markers to assess liver fibrosis severity [14]. High OPG expression was also observed in mouse models of silica [15] and bleomycin-induced pulmonary fibrosis [16]. OPG levels increase soon after bleomycin treatment and wane over time, accompanied by a decrease in deposited collagen. We therefore reasoned that OPG expression may be a marker of active fibrotic processes in human pulmonary fibrosis and may also be used as a marker to diagnose pulmonary fibrosis and/or track or predict progression. We investigated this hypothesis using mouse and human lung tissue and serum of patients with IPF and control individuals.

## MATERIALS AND METHODS

### Human tissue

Human fibrotic lung tissue was collected with informed consent from patients with end-stage IPF at either the University Medical Centre Groningen (UMCG) or at the Erasmus Medical Centre Rotterdam. Nonfibrotic control lung tissue was obtained at the UMCG from patients undergoing surgical resection for carcinoma or chronic obstructive pulmonary disease (COPD, patients characteristics are summarized in table 1).

**Table 1.**
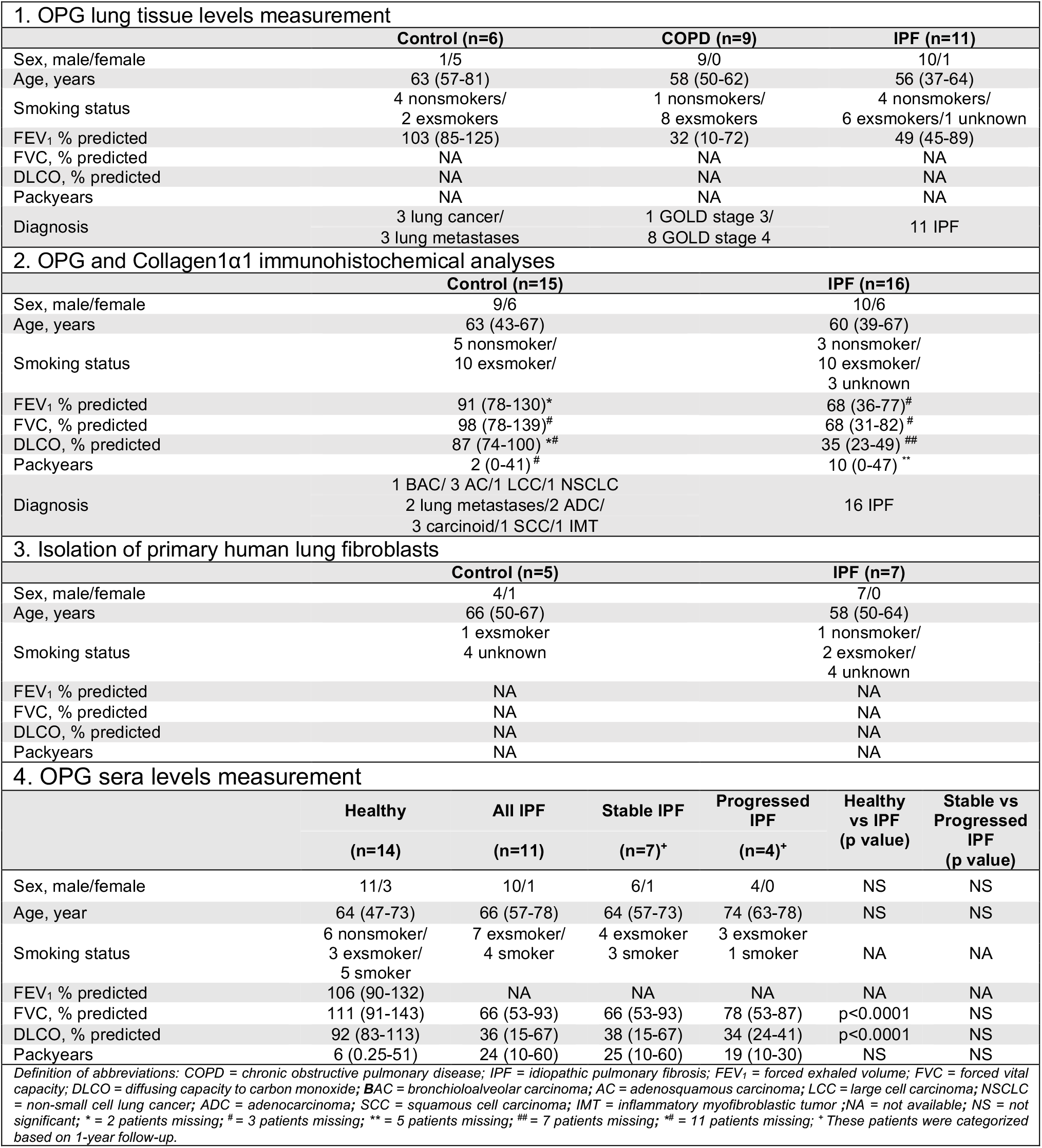
Characteristics of patients whose lung tissue or sera were used for OPG or Collagen1α1 analyses and fibroblast isolation (medians with range are presented).

### Isolation and culture of primary fibroblasts

Primary human lung fibroblasts were isolated from 5 non-fibrotic control patients and 7 patients with IPF (patient characteristics are summarized in table 1). Plated cells were grown to confluence and transferred to low-serum medium. After 24h, supernatants were collected and stored at -20 °C until further analyses.

### Serum from patients with IPF and healthy controls

Sera from 11 patients with IPF were collected in tertiary care centre for ILD, University of Heidelberg, Germany (IRB vote number S-270/2001). Diagnosis of IPF was made in a multidisciplinary team according to current international guidelines [17, 18] Sera from 14 matched healthy individuals were collected at the UMCG, Netherlands (METC numbers 2009/007 and 2016/572, detailed characteristics of the patients and healthy individuals are summarized in table 1). Patients with IPF were followed for a minimum of 1 year after first presentation. Patients were categorized as having progressed if, within a 1-year monitoring period, the percentage predicted diffusing capacity of the lung for carbon monoxide (DLCO) decreased more than 15% or the percentage predicted force vital capacity (FVC) decreased more than 10%, or in case of death [19].

### Animal experiments

All animal experiments were approved by the Institutional Animal Care and Use Committee, The University of Groningen (DEC6064) and (DEC6416AA). Male C57BL/6 mice were obtained from Harlan (Zeist, The Netherlands). Fibrosis was induced using intratracheal installation of Min-U-Sil 5 crystalline silica (0.2 g/kg) [15]. Mice were sacrificed after 6 weeks and serum and lung tissues collected.

Lungs of healthy male C57BL/6 mice were used to make precision-cut lung slices. Slices were incubated for 48h with or without 5 ng/ml TGFβ1 in the presence or absence of 10 μM galunisertib. Slices and supernatants were collected for further analyses.

### Protein isolation from lung tissue

Protein was isolated from frozen human or mouse lung tissue in 0.25 M Tris/HCl buffer with 2.5% Igepal, 0.5% SDS, Protease Inhibitor Cocktail (Boehringer Mannheim, Germany), pH 7.5. Samples were stored at -80°C for later analyses.

### ELISA

Human and mouse OPG levels in lung tissue, serum and culture supernatants were assessed using an ELISA kit according to manufacturer’s instructions (R&D, Minneapolis, USA)

### Immunohistochemistry

Immunohistochemical analyses of collagen I, collagen 1α1 (Col1α1) and OPG were performed on paraffin sections of mouse or human lung tissue. The amount of collagen I deposition in the mouse lung was quantified using ImageScope software (Aperio, Burlingame, USA) and OPG and Col1α1 in human lung were assessed using ImageJ software (version 1.47, NIH, USA) [20].

### Quantitative Real-time PCR

Total mRNA was isolated from tissues using a Maxwell^®^ LEV simply RNA Cells/Tissue kit (cat# AS1280, Promega, Madison, WI) according to the manufacturer’s instructions. Col1α1, fibronectin, OPG, plasminogen activator inhibitor-1 (PAI1), and 18s mRNA was detected with primers summarized in supplemental table 1. mRNA expression was normalized to 18s.

### Statistical analyses

Statistical differences between two groups without normal distribution were assessed using Mann-Whitney U for unpaired data or Wilcoxon for paired data. D’agostino-Pearsons test was used to examine normality. Paired or unpaired student’s t-test was used for normal data. For the comparison of multiple groups, we used Kruskal Wallis. Correlations were assessed by Spearman or Pearson test. Receiver-operator characteristic (ROC) curves and Kaplan-Meier curves were used to model the utility of serum OPG as a marker of disease progression and to model survival respectively. Significance was considered when p<0.05. The data were analysed using GraphPad Prism 8 (GraphPad software, San Diego, USA).

More detailed information is available in the online supplemental “Material and Methods” section.

## RESULTS

### OPG protein levels are higher in mice with silica-induced lung fibrosis and in human fibrotic lung tissues

We first examined OPG levels in the lungs of silica-treated mice and observed that OPG levels were significantly higher in fibrotic lungs compared to control lungs (fig. 1a). To confirm the development of fibrosis in our mouse model we examined collagen I protein deposition in lungs of silica-exposed mice and found significantly higher collagen expression and deposition in lungs of mice exposed to silica (supplemental data fig. 1a-b). We also measured OPG levels in serum, but found no significant differences between control and silica-exposed mice (supplemental data fig. 1c). We then measured OPG levels in lung tissue from patients with end stage IPF and found that OPG levels were significantly higher when compared to lung tissue of control patients (Figure 1b).

**Figure 1.**
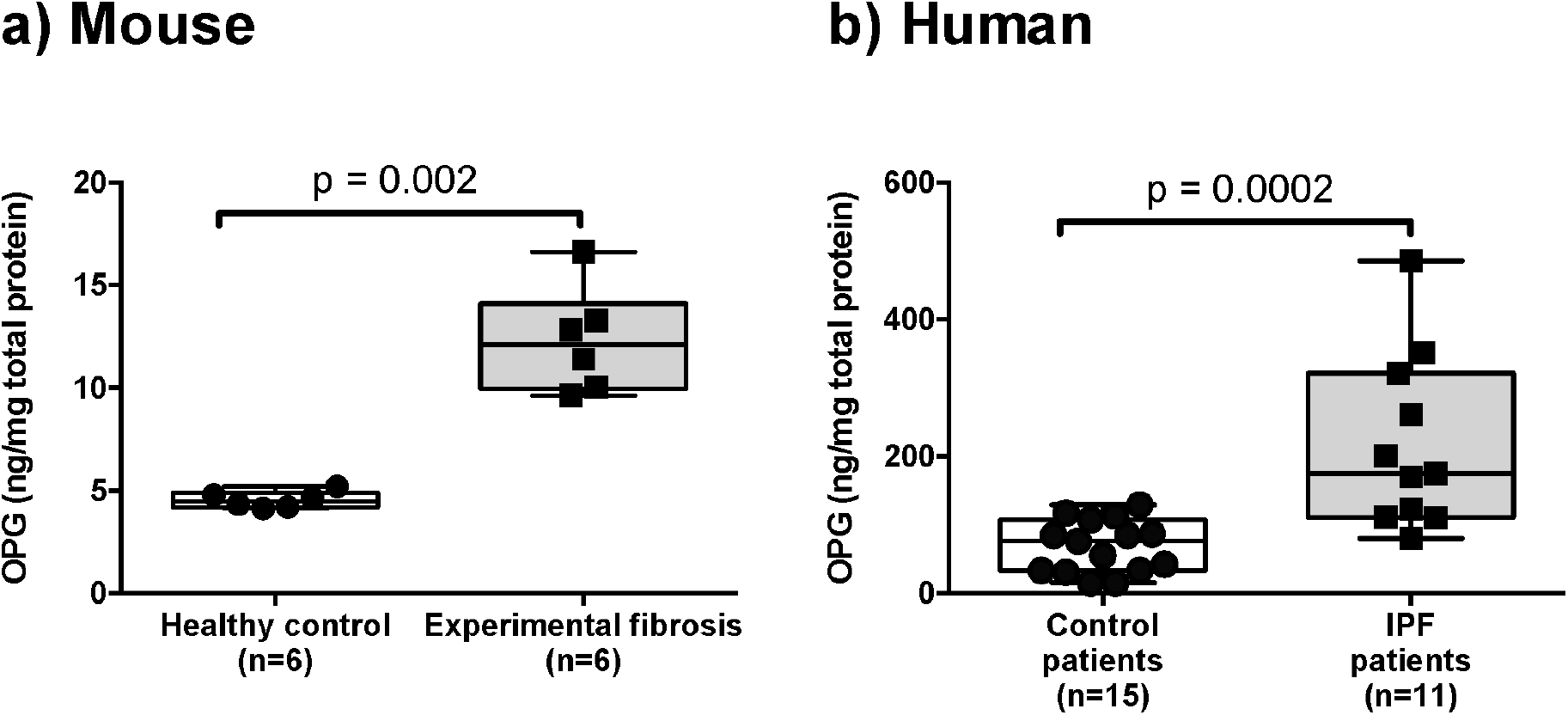
OPG protein levels are higher in fibrotic conditions in both mouse and human lung tissue. (a) OPG protein levels were measured in mouse lung tissues collected from mice 6 weeks after receiving silica (experimental fibrosis, n=6) or saline (healthy control, n=6). (b) in human lung tissue obtained from 15 patients undergoing surgical resection for carcinoma or COPD (control patients) and from 11 patients with idiopathic pulmonary fibrosis (IPF) undergoing lung transplantation. Differences between groups were tested with Mann Whitney U test (mouse) or unpaired student t test (human), p<0.05 was considered significant.

### TGFβ induces expression of OPG and fibrosis-associated markers in mouse precision-cut lung slices

To determine whether OPG can be produced by lung tissue, we added the profibrotic cytokine TGFβ1 to precision-cut lung slices of healthy mouse lung tissue. To further investigate the association between OPG production and fibrosis development and the potential regulation by antifibrotic treatment, we conducted these experiments with or without galunisertib, a TGFβ-receptor type I kinase inhibitor. We found significantly higher mRNA expression and protein production of OPG, along with the fibrosis-associated genes PAI1 and fibronectin following TGFβ1 treatment as compared to untreated controls and this could be blocked by galunisertib (fig. 2a-c). Although mRNA expression of Col1α1 (fig. 2d) was not significantly higher after TGFβ1 treatment, galunisertib still significantly attenuated TGFβ1-induced levels of Col1a1 mRNA.

**Figure 2.**
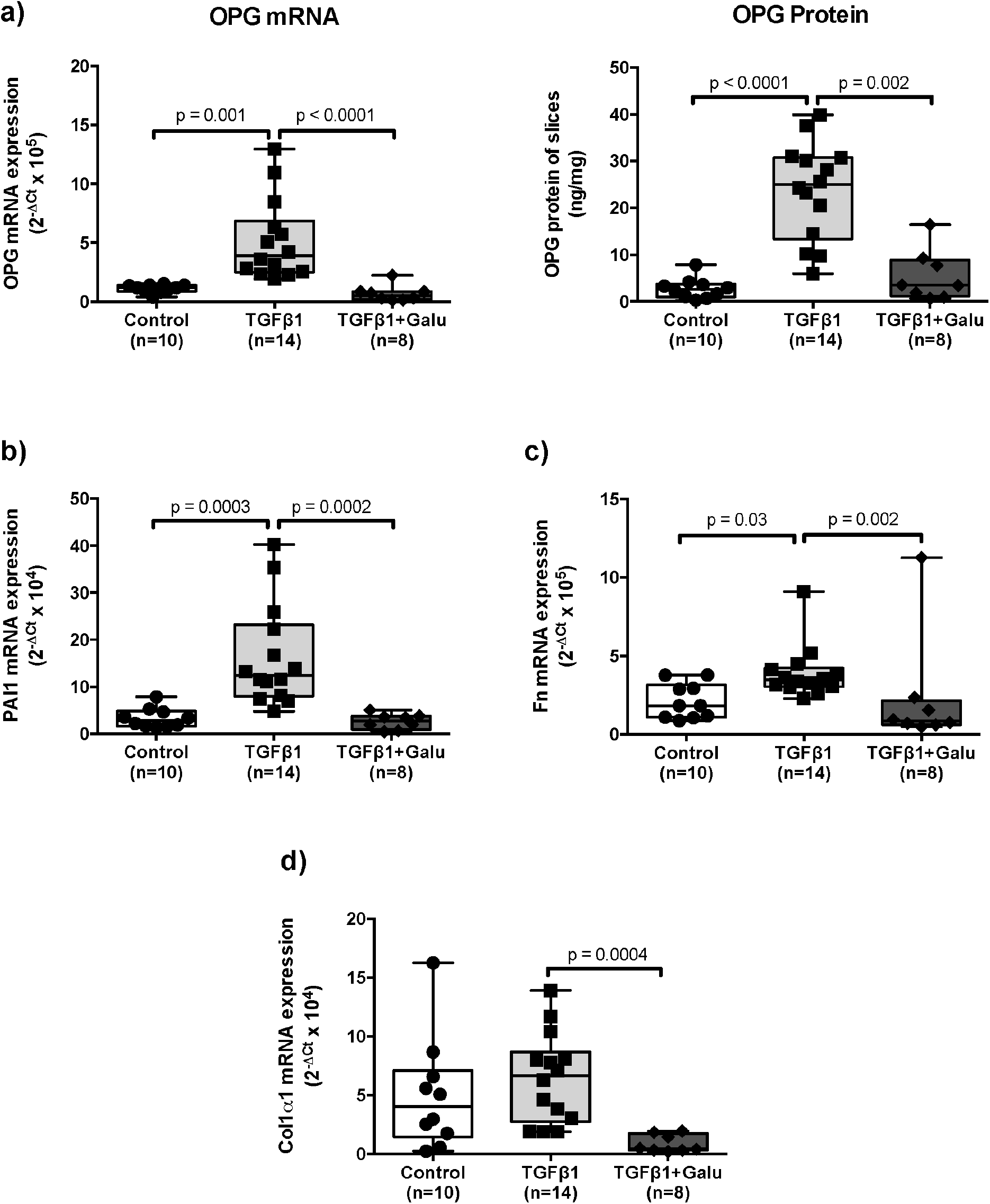
TGFβ induces expression of OPG and fibrosis-associated markers in mouse precision-cut lung slices. Mouse lung slices were treated with 5 ng/mL TGFβ1 with or without 10 μM galunisertib for 48 hours and compared to untreated controls for expression of (a) OPG mRNA and OPG protein, fibrosis-associated markers (b) plasminogen activator inhibitor-1 (PAI1), (c) fibronectin (Fn), and (d) Col1α1 mRNA. Differences between groups were analysed with a Kruskal Wallis test. p<0.05 was considered statistically significant.

### OPG expression correlates with fibrosis-associated markers in mouse lung slices

To investigate whether the expression of OPG correlated with fibrosis-associated markers, we compared OPG mRNA expression levels with those of Col1α1, fibronectin, and PAI1 in all treatment groups of slices (fig. 2). We found that OPG mRNA expression significantly correlated with mRNA expression of Col1α1 (fig. 3a), fibronectin (fig. 3b) and PAI1 (fig. 3c).

**Figure 3.**
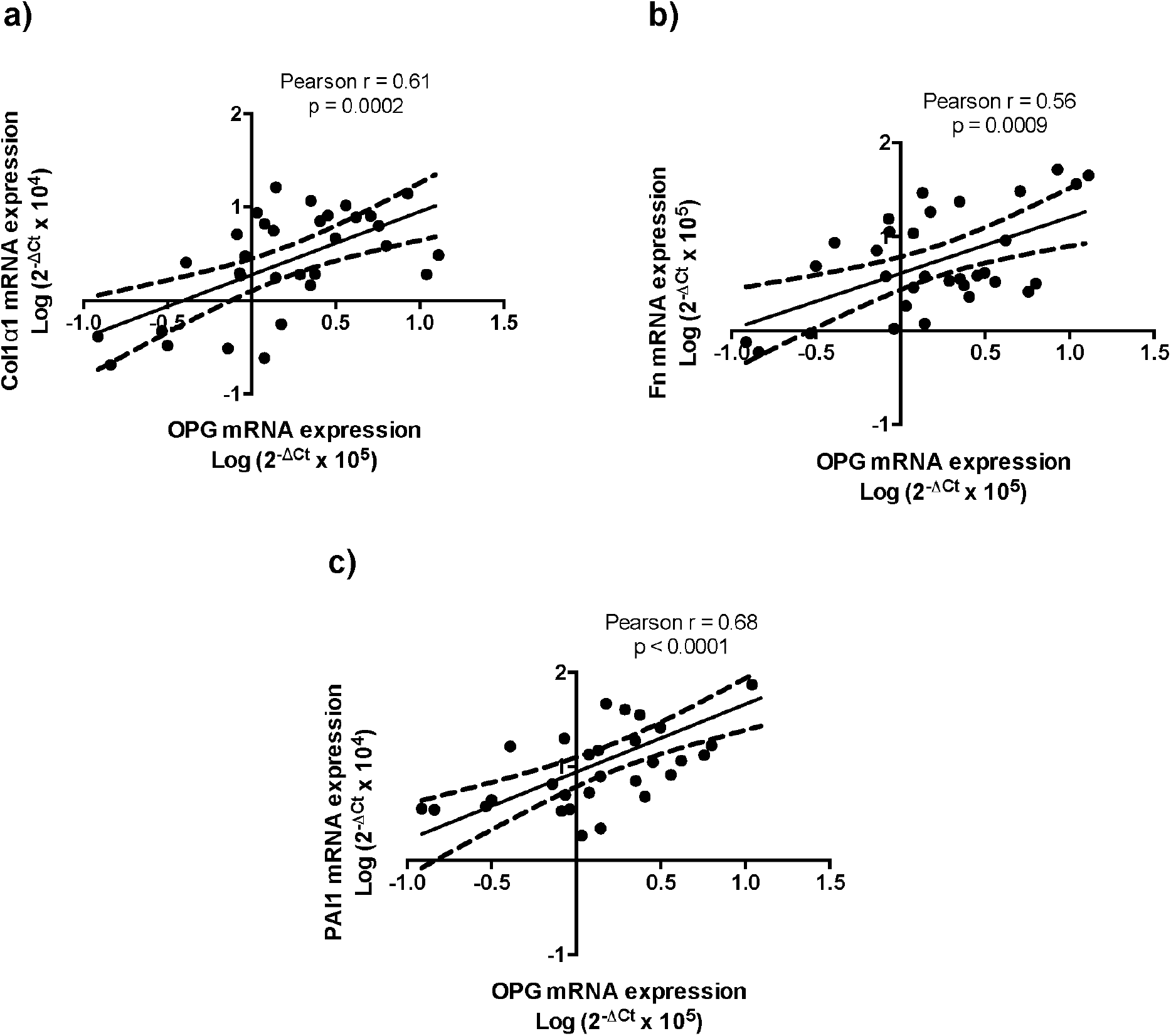
OPG mRNA expression levels correlate with expression of fibrosis-associated markers in mouse lung slices. In untreated mouse lung slices and slices treated with TGFβ1 with or without galunisertib, OPG mRNA expression correlated positively with mRNA expression of fibrosis-associated markers (a) Col1α1, (b) fibronectin (Fn) and (c) PAI1. Correlations were calculated using a Pearson test and p<0.05 was considered significant.

### OPG is abundantly present in fibrotic areas in mouse lung tissue and OPG protein levels positively correlate with the degree of fibrosis

To investigate whether OPG expression could be related to the degree of fibrosis in lung tissue, we assessed OPG (fig. 4) and collagen I deposition (supplemental data fig. 1a-c) in lung tissue of mice exposed to silica. Immunohistochemistry for OPG showed that OPG localized in the smooth muscle layers around vessels and airways in lung tissue of control mice (fig. 4a). In lung tissue of mice exposed to silica, OPG expression was also observed within areas of fibrosis (fig. 4b). Furthermore, we found that OPG protein expression significantly correlated with collagen I deposition in lung tissue of mice (Fig. 4d). OPG presence in mouse lung tissue did not correlate with OPG levels in serum (supplemental data fig. 2a). and neither did serum OPG levels correlate with collagen I content in lung tissue (supplemental data fig. 2b)

**Figure 4.**
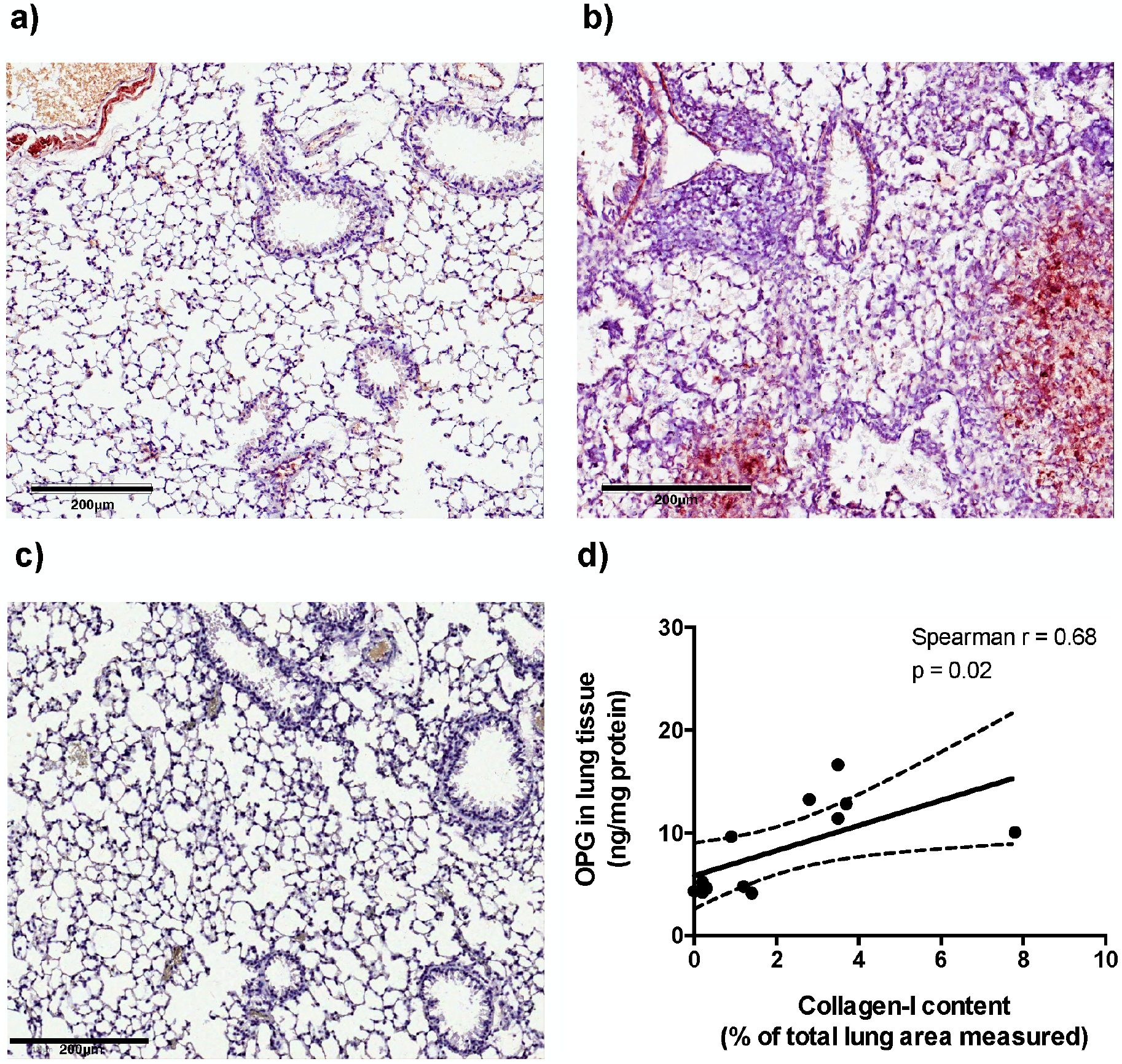
OPG is abundantly present in fibrotic areas in lung tissue and OPG protein levels positively correlate with the degree of fibrosis in mice. Mice were exposed to silica or saline and sacrificed after 6 weeks. Representative sections of lung tissue stained for OPG (OPG: red, nuclei: blue) from (a) saline-exposed control and (b) silica-exposed mice. (c) Negative control for the OPG staining. (d) In lung tissue of saline-exposed control mice (n=6) and silica-exposed mice (n=6), OPG levels positively correlated with collagen I levels in lung tissue. Correlation was calculated using a Spearman test and p<0.05 was considered significant.

### OPG is abundantly present in human IPF lung tissue and is produced by lung fibroblasts

We then investigated OPG and collagen 1α1 deposition in lung tissue of patients with or without IPF. Similar to what we observed in mouse lung tissue, OPG localized to the smooth muscle layers around vessels and airways in lung tissue of control patients and was also found in epithelial cells, macrophages, and endothelial cells. In lung tissue of patients with IPF, OPG expression was additionally observed within areas of fibrosis development/fibroblast foci (fig. 5a). Furthermore, we quantified expression of OPG and found significantly more OPG-positive pixels and more intense OPG staining (Fig. 5b) in lung tissue of patients with IPF than in that of control patients. The same was found for collagen 1α1 deposition: significantly more collagen 1α1-positive pixels and more intense collagen 1α1 staining (Fig. 5c) in lung tissue of patients with IPF than in lung tissue of control patients. Despite these similarities in staining, we did not find a significant correlation between OPG and collagen 1α1 deposition in human IPF lung tissue as we had seen in mice (data not shown). For both types of staining we found a lower percentage of positively stained area in IPF lung tissue compared to control, due to the increase in tissue mass in fibrotic lungs compared to control lung tissues (supplemental data fig. 3a-b).

**Figure 5.**
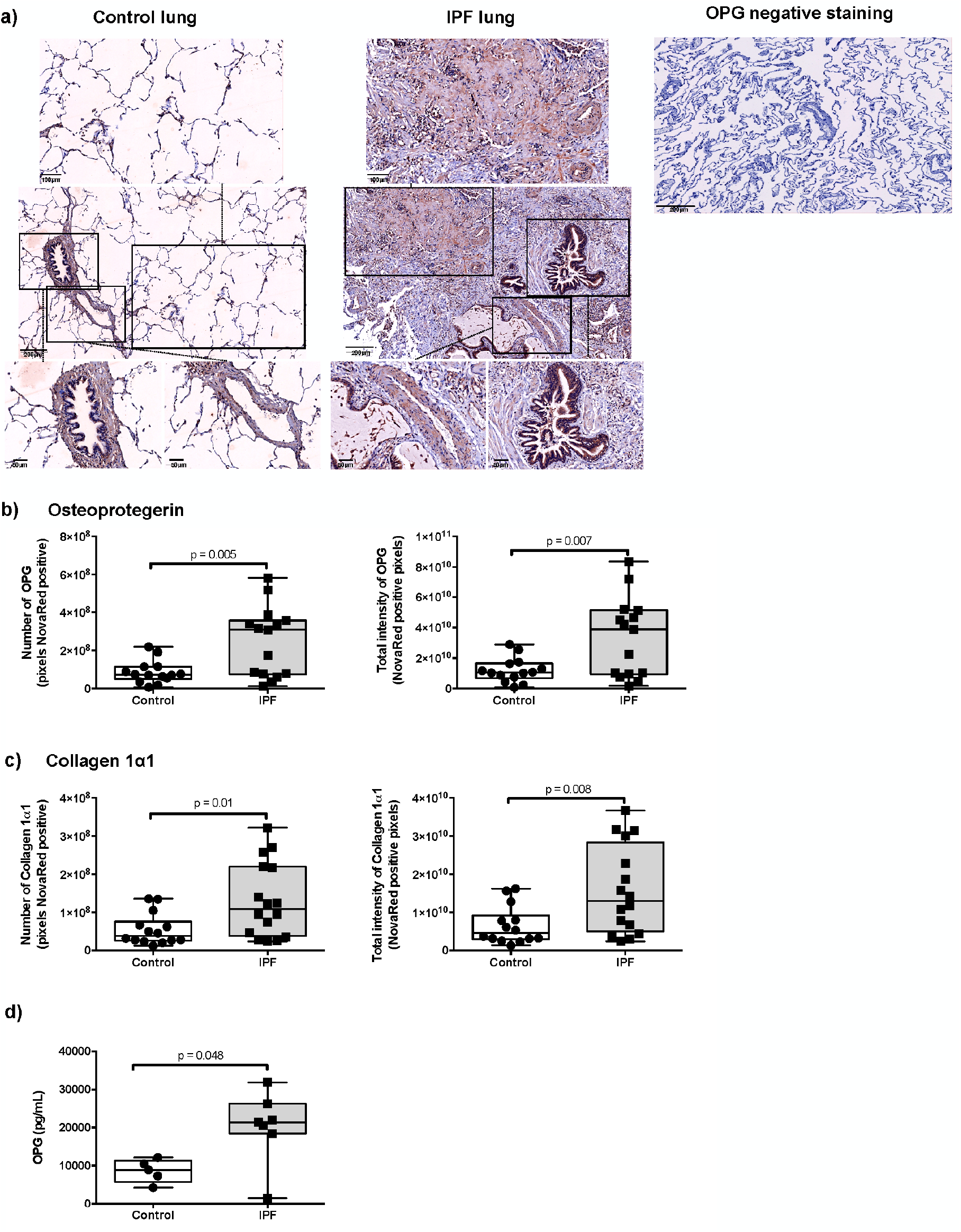
OPG is abundantly present in human IPF lung tissue and produced by lung fibroblasts. (a). Representative sections of lung tissue stained for OPG (OPG; red, nuclei; blue) from a control patient, a patient with IPF and negative control for the OPG staining. (b) Numbers of OPG-positive pixels and total intensity of OPG-positive pixels, (c) numbers of collagen 1α1-positive pixels and total intensity of collagen 1α1-positive pixels were quantified using imageJ. (d) Human primary fibroblasts isolated from lung tissue of nonfibrotic control patients and patients with IPF were cultured for 24 hours and levels of secreted OPG protein were quantified. Differences between groups were tested with unpaired student t test (b-c) or Mann Whitney U test (d). p<0.05 was considered significant.

As more OPG staining was observed in fibrotic area/fibroblast foci, we investigated if (myo)fibroblasts could be a source of OPG. To this end, we isolated primary human fibroblasts from patients with IPF and from control patients. We observed that human lung fibroblasts produced copious amounts of OPG and that lung fibroblasts from patients with IPF produced significantly more OPG than lung fibroblasts isolated from control patients (fig. 5d).

### Serum OPG levels negatively correlate with DLCO and associate with disease progression in patients with IPF

To determine whether OPG could serve as a blood-based biomarker for IPF, we measured OPG levels in serum from patients with IPF, collected at first presentation and compared them to serum of age and gender matched healthy individuals. Serum OPG levels of patients with IPF were not significantly different as compared to healthy individuals (fig. 6a). However, patients with IPF had more variable OPG levels than healthy controls, similar to the pattern that was seen for OPG levels in lung tissue (fig. 1a). To investigate whether OPG levels reflect disease status, we examined correlations between serum OPG levels and lung function parameters in patients with IPF at first presentation and found an inverse correlation of elevated OPG levels to decreased DLCO levels (Pearson r=-0.64, p=0.035) (fig. 6b). A 1-year follow-up showed that OPG levels did not change significantly over time (supplemental data fig. 4), however, there was significant disease progression in 4 patients while 7 patients remained stable (table 1). For OPG, the area under the curve of the ROC analysis was 0.85 (95%CI, 0.62-1.09; P=0.058) (fig. 6c). Thresholds for OPG were generated from this curve and sensitivity and specificity at every OPG level calculated. At a threshold of 1234 pg/mL, serum OPG level identified patients with IPF with significant disease progression with the highest value for both sensitivity and specificity (100% and 71.43% respectively) (supplemental table 2). Finally, Kaplan-Meier analysis showed that serum OPG levels greater than 1234 pg/mL were associated with a shorter survival time than levels below 1234 pg/mL (p=0.03, fig. 6d) in this small group of patients.

**Figure 6.**
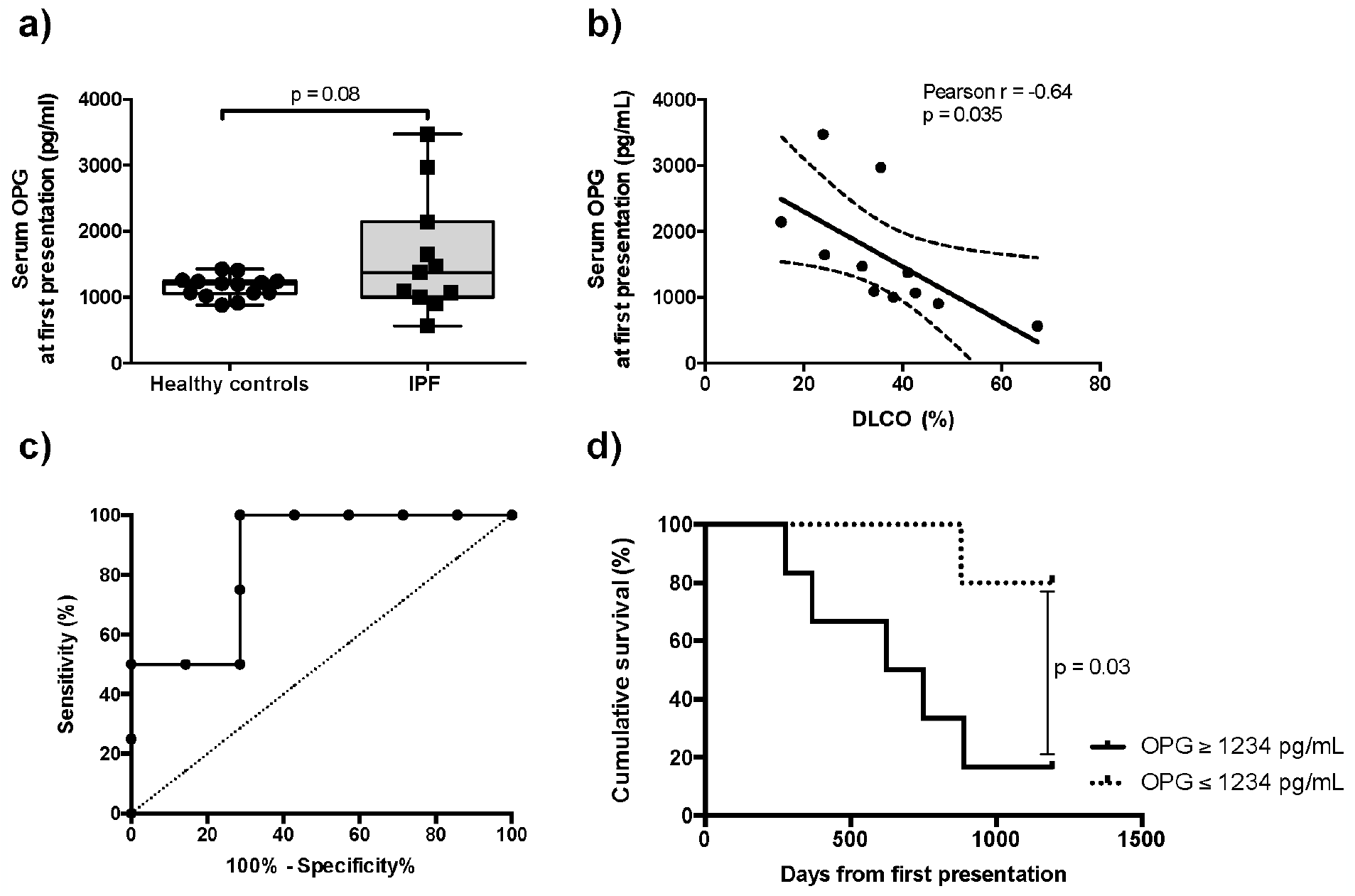
OPG levels negatively correlate with DLCO and associate with progression of disease in patients with IPF. (a) Patients with IPF were monitored for at least 1 year after the initial visit. Serum levels of OPG were quantified in blood of healthy controls and patients with IPF. (b) Serum OPG at first presentation in the hospital correlated negatively to the diffusion capacity for carbon monoxide (DLCO) in patients with IPF. (c) Receiver operating characteristic (ROC) curve was generated based on OPG levels in patients who progressed (n=4) compared to the levels of non-progressed patients (n=7, area under curve, 0.85; 95%CI, 0.62-1.09; p=0.058). (d) Survival time of patients with serum OPG levels greater than 1234 pg/mL (n=6) were compared to patients with levels below 1234 pg/mL (n=5) using Kaplan-Meier analysis. Differences between groups were tested with an unpaired t test. Correlations were calculated using a Pearson test. p<0.05 was considered significant.

## DISCUSSION

Our study has shown that OPG is expressed in the lung and is higher in fibrotic lung tissue. Interestingly, although OPG serum levels did not distinguish patients with IPF from healthy controls, elevated OPG levels at first hospital presentation were associated with disease progression. In our small cohort, patients with serum OPG levels greater than 1234 pg/mL progressed more rapidly than patients with lower levels. This is an important initial finding since information on the likelihood of disease progression is essential as it can guide clinicians to tailor clinical management of these patients. However, further validation in a larger cohort is warranted to explore the potential of OPG for assessing the prognosis of fibrotic lung disease. This finding also suggests OPG has a pathological role in IPF and that it may be important during phases or in regions of active fibrosis in the lung. Individual OPG levels in both progressed and stable patients did not change significantly over time. This may suggest that the maximal OPG production associated with an active disease process had already been attained before the patients presented at the clinic. Our studies with murine precision-cut lung slices confirm this rapid and early response of OPG to a fibrotic trigger.

Some publications have suggested that OPG levels increase in fibrotic conditions to counteract higher RANKL levels, possibly originating from bone in response to the actions of IL-6 and TNF-α [21, 22]. However, we found RANKL levels were similar to controls in patients with IPF and mice with pulmonary fibrosis (data not shown). TGFβ, the hallmark cytokine of fibrosis, induced OPG production in mouse precision-cut lung slices which could be inhibited by galunisertib. Together these data suggest that OPG in fibrotic conditions is not produced in response to RANKL but in response to TGFβ and/or possibly other profibrotic mediators. In mice, OPG levels in fibrotic lung tissue positively correlated with degree of fibrosis in lung tissue, but serum OPG levels did not correlate with the degree of fibrosis (or with lung tissue OPG levels). Interestingly, a recent study by Tsukasaki *et al*. suggested that locally produced OPG was crucial for its local functions and that its role was limited to the tissue in which it was produced [23, 24]. Moreover, we also found that lung fibroblasts isolated from IPF lung tissue produced more OPG than those from non-diseased controls. Therefore, our findings strongly support the concept that OPG is produced locally in lung tissue and is associated with the local fibrotic process.

The biological role of OPG is still under investigation. As the decoy receptor for RANKL and TRAIL, OPG could exert its effects through either of these ligands. No biological role for RANKL in lung tissue has been described, but TRAIL was shown to have anti-inflammatory effects by accelerating neutrophil apoptosis and antifibrotic effects by inducing apoptosis in (myo)fibroblasts [25-28]. Indeed, TRAIL was present at lower levels in patients with IPF and mice with bleomycin-induced pulmonary fibrosis [29]. By neutralizing TRAIL, OPG could dampen these beneficial effects, thereby contributing to progression of fibrosis.

Like McGrath *et al*. before us, we found no significant differences in serum levels of OPG between controls and patients with IPF [29] or between mice with and without fibrosis, making the use of OPG alone as a serum marker for diagnosis of IPF unlikely. We did find higher OPG levels in lung tissue of both patients with IPF and mice with fibrosis (which McGrath *et al*. did not investigate), suggesting that maybe a more lung-localized form of sampling (BAL/sputum) has merits, but this needs further investigation. We did notice more variation in OPG levels in patients with IPF than in controls (in both serum and lung tissue) and therefore investigated correlations with lung function. Unlike the previous study, we found a negative correlation between OPG levels in serum and DLCO at first presentation in patients with IPF [29]. The difference between these findings may be caused by the sampling time; we analysed serum levels of patients at the time of first presentation to our tertiary referral hospital, while McGrath used samples from patients throughout their disease progression. Interestingly, in patients with stage II sarcoidosis a negative correlation between OPG levels in BALF and DLCO was recently reported [30]. Patients with stage II sarcoidosis, however, do not have fibrotic involvement (yet), which may point again at OPG possibly being an indicator of early fibrotic activity. An alternative explanation comes from clinical studies showing that higher serum OPG levels are associated with vascular calcification [31]. If these calcifications occur in the pulmonary vascular bed, this may also explain the inverse correlation between OPG and DLCO in pulmonary fibrosis patients.

Several serum biomarkers of IPF severity and progression have been studied and characterized [32]. These include matrix metalloproteinases 1 and 7 [33], surfactant proteins SP-A and SP-D [34], SP-B [35] and fibulin-1 [36]. However, there is not a single biomarker that can accurately diagnose and predict likelihood of progression of IPF. Numerous factors are known to contribute to fibrosis development and these may vary between patients. Therefore, a panel of biomarkers may be a more useful approach and OPG may contribute important information to this panel. An excellent example is seen in the liver field, in which a panel of serum markers for staging of liver fibrosis severity performed better with the addition of OPG [37].

In summary, our study revealed that OPG expression is upregulated in pulmonary fibrosis, is induced by TGFβ, and corresponds to the degree of fibrosis in the lungs of mice. We translated our findings to patients with IPF: serum OPG levels negatively correlated with DLCO and above a threshold of 1234pg/ml was associated with disease progression. This suggests possible value of OPG as a prognostic marker for likelihood of disease progression and warrants evaluation in a larger cohort of patients with IPF and other fibrotic lung diseases.

## Author contributions

Habibie: Study design, collection and assembly of data, data analysis and interpretation, manuscript writing, critical reading and revision.

Kurnia S.S Putri: Collection and assembly of data, data analysis and interpretation, critical reading and revision.

Carian E. Boorsma: Collection and assembly of data, data analysis and interpretation, critical reading and revision.

David M. Brass: Study design, critical reading and revision.

Peter Heukels: Collection and assembly of data, experimental material support, critical reading and revision.

Marlies Wijsenbeek: Collection and assembly of data, experimental material support, critical reading and revision.

Mirjam Kool: Collection and assembly of data, experimental material support, critical reading and revision.

Maarten van den Berge: Collection and assembly of data, experimental material support, critical reading and revision.

Theo Borghuis: Collection and assembly of data, data analysis and interpretation, critical reading and revision.

Corry-Anke Brandsma: Collection and assembly of data, experimental material support, critical reading and revision.

C Tji Gan: Collection and assembly of data, experimental material support, critical reading and revision.

Peter Olinga: Collection and assembly of data, experimental material support, critical reading and revision.

Wim Timens: Collection and assembly of data, experimental material support, critical reading and revision.

Nicolas Kahn: Collection and assembly of data, experimental material support, data analysis and interpretation, critical reading and revision.

Michael Kreuter: Collection and assembly of data, experimental material support, data analysis and interpretation, critical reading and revision.

Janette K. Burgess: Study design, data analysis and interpretation, manuscript writing, critical reading and revision.

Barbro N. Melgert: Collection and assembly of data, study design, data analysis and interpretation, financial support, manuscript writing, critical reading and revision.

## SUPPLEMENTAL DATA

## MATERIALS AND METHODS

### Human tissue

Human fibrotic lung tissue was collected with informed consent from patients with end-stage IPF undergoing lung transplantation at either the University Medical Centre Groningen (UMCG) or at the Erasmus Medical Centre Rotterdam. Human nonfibrotic control lung tissue was obtained at the UMCG from patients undergoing surgical resection for carcinoma or chronic obstructive pulmonary disease (COPD). In cases of tumour resections, histologically normal lung tissue was taken as far distally as possible from the tumour and assessed visually by a pathologist for abnormalities with a standard haematoxylin and eosin staining. Frozen lung tissue was used for the analysis of protein levels of OPG and formalin-fixed paraffin embedded lung tissue was used for the immunohistochemical analyses of OPG and collagen 1α1.

In Groningen, the study protocol was consistent with the Research Code of the University Medical Centre Groningen, (www.umcg.nl/EN/Research/Researchers/General/ResearchCode/Paginas/default.aspx) and the Dutch national ethical and professional guidelines (“Code of conduct; Dutch federation of biomedical scientific societies”; http://www.federa.org). In Rotterdam, the Medical Ethical Committee approved all protocols followed in that centre.

### Isolation of primary human lung fibroblasts

Primary human lung fibroblasts were isolated from lung tissue obtained from 5 nonfibrotic control patients and 7 patients with IPF using a technique described previously [1, 2]. Parenchymal human lung tissue, excluding visible vessels and airways, was cut into 1-2 mm^2^ pieces that were then cultured for 4-5 weeks in 12-well plates in the presence of complete Ham’s F12 medium (BioWhittaker Europe BV, Belgium) supplemented with 10% fetal calf serum (FCS, Invitrogen, The Netherlands), L-glutamin 2mM, fungizone, penicillin 100 U/mL and streptomycin 100 µg/mL at 37°C in an atmosphere of 5% CO_2_. At 25% confluency, the cells were transferred to T25 flasks. At confluence, cells were trypsinized and cryopreserved in FCS with 10% DMSO under slow cooling conditions and eventually stored at -150°C. Cells used in the experiments described were maximally passaged seven times.

### OPG measurement from primary human lung fibroblasts

Cells were seeded in 96-wells plates (3200 cells/well) for experiments and maintained in Dulbecco’s Modified Eagle’s Medium (DMEM) Low Glucose (Biowest, Nuaillé, France) supplemented with 10% Fetal Bovine Serum (FBS), glutaMAX, penicillin (100 U/mL) and streptomycin (100 µg/mL). Plated cells were grown for 24h and transferred to low-serum medium DMEM low glucose supplemented with 0.1% BSA (Sigma-Aldrich, St. Louis, MO, USA) glutaMAX, penicillin (100 U/mL) and streptomycin (100 µg/mL). After 24h, supernatants were collected for ELISA analysis of OPG levels and stored at -20°C for later analysis.

#### Serum from IPF patients and healthy controls

Sera were collected, with informed consent, from 11 IPF patients at their first presentation at the German Centre for Lung Research, University of Heidelberg, Heidelberg, Germany (IRB vote number S-270/2001) and these were compared to sera from 14 matched healthy individuals collected at the UMCG, Groningen, Netherlands (METC numbers 2009/007 and 2016/572). IPF diagnosis was assessed according to current guidelines by a multidisciplinary team.

#### Animal experiments

Male C57BL/6 mice were obtained from Harlan (Zeist, The Netherlands). Animals were maintained with permanent access to food and water in a temperature-controlled environment with a 12h dark/light cycle regimen in groups of three with cage enrichment. Treatment with silica did not result in implementing humane end points to prevent further distress and all 12 animals were used for analysis. All animal experiments were approved by the Institutional Animal Care and Use Committee. Mice were used in experiments with silica-induced pulmonary fibrosis (DEC6064) and for the preparation of precision cut lung slices (DEC6416AA). Animal experiments were performed in the animal facility of the University of Groningen according to strict national and international guidelines on animal experimentation.

#### Silica-induced pulmonary fibrosis in mice

Male C57BL/6 mice (20-30 gr, age 8-12 weeks, n=12) were randomly selected to serve as controls (n=6) or to receive silica to induce fibrosis (n=6) and mice were housed in randomly established groups to avoid cage effects. Sample size was calculated based on pilot experiments. Fibrosis was induced using intratracheal installation of Min-U-Sil 5 crystalline silica (a kind gift from Dr. Andy Ghio, US EPA, Chapel Hill, NC). Mice were anesthetized using isoflurane before they received a single administration of crystalline silica (0.2 g/kg) in 50 μl 0.9% saline by intratracheal installation, while control animals received an equivalent volume of 0.9% saline. Mice were sacrificed after 6 weeks and serum and lung tissue were collected for OPG levels and collagen I content measurements.

#### Mouse precision-cut lung slices

An *ex vivo* model of early pulmonary fibrosis development was used to investigate regulation of OPG production by lung tissue. Lungs of 14 healthy male C57BL/6 mice (20-30 gr, age 8-12 weeks) were used to make precision-cut lung slices using a method described previously [3]. After sacrifice by exsanguination via the aorta abdominalis under isoflurane anaesthesia, mouse lungs were filled with 1.5% low-melting temperature agarose in 0.9% NaCl (Sigma-Aldrich) and transferred directly into ice-cold University of Wisconsin organ preservation solution (DuPont Critical Care, Waukegab, IL). Lung slices of 5-mm diameter and a weight of about 5 mg (or thickness of 250-300 µm), were prepared with a Krumdieck tissue slicer (Alabama Research and Development, AL) using ice-cold Krebs-Henseleit Buffer [25 mM D-glucose (Merck, Darmstadt, Germany), 25 mM NaHCO_3_ (Merck), 10 mM HEPES (MP Biomedicals, Aurora, OH), saturated with carbogen (95% O_2_/5% CO_2_) and adjusted to pH 7.4]. Lung slices were incubated in 12-well plates in 1.3 mL DMEM + Glutamax medium [4.5g/L D-glucose and pyruvate (Gibco) supplemented with a non-essential amino acid mixture (1:100), 100 U/ml penicillin, 100 μg/ml streptomycin, 45 µg/ml gentamycin and 10% FCS]. After a 1h pre-incubation at 37°C in a 80% O_2_/5% CO_2_ atmosphere with continuous shaking at 90 rpm, slices were transferred into fresh medium and incubated for 48h with or without TGFβ (5 ng/ml) to induce early fibrosis in the presence or absence of galunisertib (10µM). Medium and treatments were refreshed after 24 hours. Four slices per treatment were pooled for further analyses, while supernatants were collected separately. All samples were stored at -80°C for further analyses.

#### Protein isolation from lung tissue

Protein was isolated from frozen human or mouse lung tissue (30-40 mg) in 0.25 M Tris/HCl buffer, 2.5% Igepal, 0.5% SDS, Protease Inhibitor Cocktail (Boehringer Mannheim, Germany), pH 7.5 by incubating for 1 hour at 4°C, homogenizing and centrifuging at 16,000 x g for 30 min. Supernatants were collected and the overall protein concentration of the tissue lysates were determined using an RC DC™ Protein Assay based on the Lowry method (cat#5000111, Bio-Rad, Hercules, CA). Samples were stored at -80°C until further analysis by ELISA analysis.

#### ELISA

Human and mouse OPG levels in lung tissue, serum and culture supernatants were assessed using ELISA (cat#DY805 (human), cat#DY459 (mouse), R&D Systems (Minneapolis, MN, USA) according to the instructions provided by the manufacturer. One hundred micrograms of tissue lysate total protein in a total of 100 μl of sample buffer were analysed. Cell culture media, lung slice culture medium and serum were diluted 1:10, 1:5 and 1:1 respectively before analysis. Total OPG in the lung tissue and lung slices culture medium was corrected for the protein content of the tissue or slices, which was measured by Lowry (BIO-rad RC DC Protein Assay, Bio Rad, Veenendaal, The Netherlands).

#### Immunohistochemistry

Immunohistochemical analysis of collagen I and OPG protein in mouse lung tissue was performed on 3 μm paraffin-embedded sections of tissue fixed with a zinc-containing buffer [4]. Tissue sections were deparaffinized in xylene, rehydrated in alcohol, rinsed in milliQ water and placed in PBS. Sections were subjected to antigen retrieval (0.1 N Tris-HCl pH 9.0 buffer, overnight at 80°C). Prior to the incubation with OPG antibody, sections were blocked with 1% BSA in 5% nonfat milk (Sigma-Aldrich, St. Louis, MO, USA). Primary antibodies were then incubated for 1h at room temperature in the presence of 5% normal mouse serum. Antibodies used were goat anti-mouse collagen I (1:75, Southern Biotech, Birmingham, AL), rabbit anti-mouse OPG (1:400, Antibodies-online, Atlanta, GA). Primary antibody incubation was followed by incubation with either a peroxidase-conjugated goat anti-rabbit (1:100, DAKO) or a rabbit anti-goat (1:100, DAKO) secondary antibody, respectively. PO-labeled antibodies were visualized using ImPACT NovaRED kit (Vector, Burlingame, USA) with a haematoxylin counterstain. The amount of collagen deposition in the mouse lung was calculated using ImageScope software (Aperio, Burlingame, USA). After selecting the stained area of the lung sections exluding edges of the tissue, large airway and vessels, a threshold was set to identify positive staining of the tissue (represented by collagen). The percentage of stained-tissue surface per total-tissue surface analysed was then calculated for each section.

Immunohistochemical analysis of collagen 1α1 and OPG protein in human lung tissue was performed on formalin-fixed paraffin sections. Human lung sections were deparaffinized and incubated with citrate for antigen retrieval at 100 °C for 15 minutes. After blocking endogenous peroxidases, slides were incubated overnight with anti-OPG (1:300, Abcam) or Col1a1 (1:500, Abcam) in PBS with 1% BSA at 4 °C overnight followed by a goat anti-rabbit peroxidase conjugated secondary antibody (1:100, DAKO) at room temperature for 1 hour. For colour development, NovaRED kit (Vector, Burlingame, USA) was applied to the slides and hematoxylin was used as a nuclear counter stain. Images were captures using a slide scanner (Nanozoomer 2.0 HT, Hamamatsu Photonics) with 20× magnification. Fiji ImageJ was used to quantify the density and distribution of staining [5]. Colour deconvolution vectors in ImageJ were optimized to ensure accurate separation of Haematoxylin and NovaRed [6]. The haematoxylin image was used to measure total tissue surface and the NovaRed image was used to calculate the number of positive pixels above the threshold within the image (‘area’) and the average intensity of the pixels above the threshold (‘average intensity’). Macros were used to batch process the images. Data were represented as percentage tissue surface area (NovaRed area/total tissue surface area) and density (average intensity calculated for the whole tissue).

#### Quantitative Real-time PCR

Total mRNA was isolated from tissues using a Maxwell^®^ LEV simply RNA Cells/Tissue kit (cat# AS1280, Promega, Madison, WI) according to the instructions provided by the manufacturer. Final mRNA concentrations were measured using a Nanodrop ND-100 spectrophotometer (Nanodrop Technologies, Wilmington, DE). All primers were obtained from Sigma-Aldrich (Zwijndrecht, The Netherlands, primers detail can be found in supplemental table 1). Conversion of RNA into cDNA was performed using a reverse transcriptase kit (Promega, Leiden, The Netherlands) in a master-cycler gradient (25 °C for 10 min, 45 °C for 60 min and 95 °C for 5 min). Transcript levels of these genes were measured by using 20 ng cDNA per sample in a SensiMix™ SYBR kit (Bioline, Taunton, MA) and an ABI7900HT sequence detection system (Applied Biosystems, Foster City, CA) with 45 cycles of 10 min 95 °C, 15 sec at 95 °C and 25 sec at 60 °C followed by a dissociation stage. For each sample, the threshold cycles (Ct values) were calculated using SDS 2.3 software (Applied Biosystems), and mRNA expression was normalized to 18s.

#### Statistical analyses

Data sets with n ≤8 were considered non-normally distributed. Statistical differences between two groups were assessed using nonparametric Mann-Whitney U for unpaired data or Wilcoxon for paired data. For data sets with n ≥8, D’agostino-Pearsons test was used to examine normality of the data. If data were normally distributed, a paired or unpaired student’s t-test was used to compare two groups depending on matched data. For the comparison of multiple groups, we used Kruskal Wallis. Correlations were assessed by calculating a nonparametric (Spearman) or parametric (Pearson) correlation coefficient (r). Receiver-operator characteristic (ROC) curves and Kaplan-Meier curves were used to model the utility of serum OPG as a marker of disease progression and to model survival respectively.

**Supplemental table 1.**
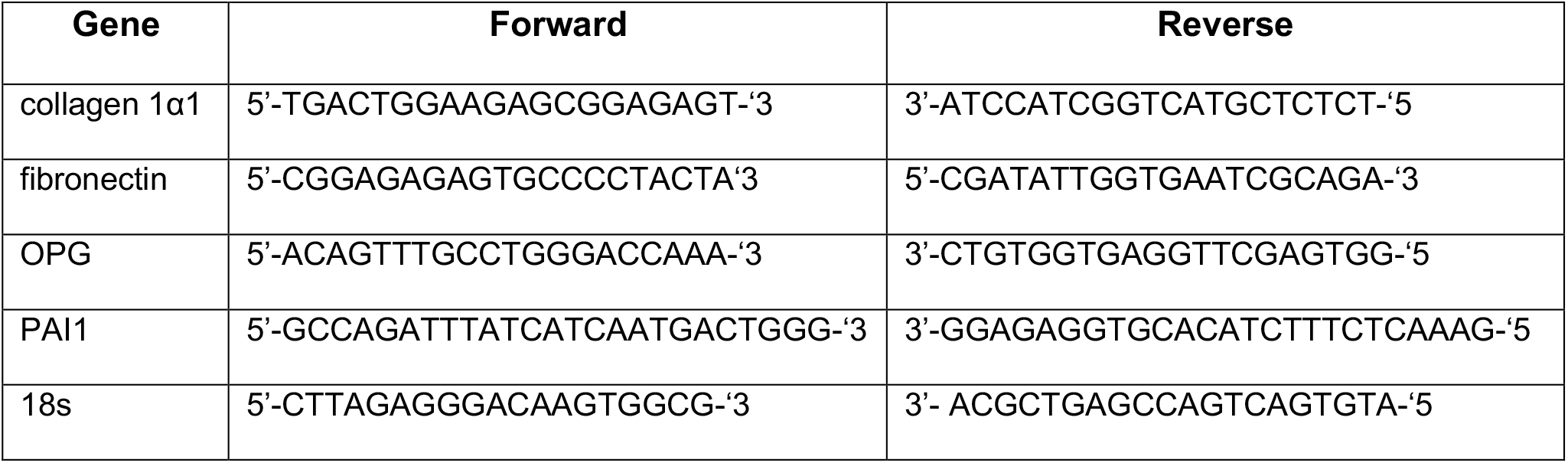
Characteristics of primers used for the quantitative real time PCR

**Supplemental table 2.** Selected threshold values from the ROC curve (area under curve, 0.85; 95%CI, 0.62-1.09; p=0.058; n=11). Values in bold were used for further analysis

| Threshold value (pg/ml) | Sensitivity% | Specificity% |
| --- | --- | --- |
| > 735.9 | 100,0 | 14,29 |
| > 954.1 | 100,0 | 28,57 |
| > 1035 | 100,0 | 42,86 |
| > 1080 | 100,0 | 57,14 |
| **> 1234** | **100,0** | **71,43** |
| > 1423 | 75,00 | 71,43 |
| > 1558 | 50,00 | 71,43 |
| > 1896 | 50,00 | 85,71 |
| > 2560 | 50,00 | 100,0 |
| > 3225 | 25,00 | 100,0 |

**Supplemental data figure 1.**
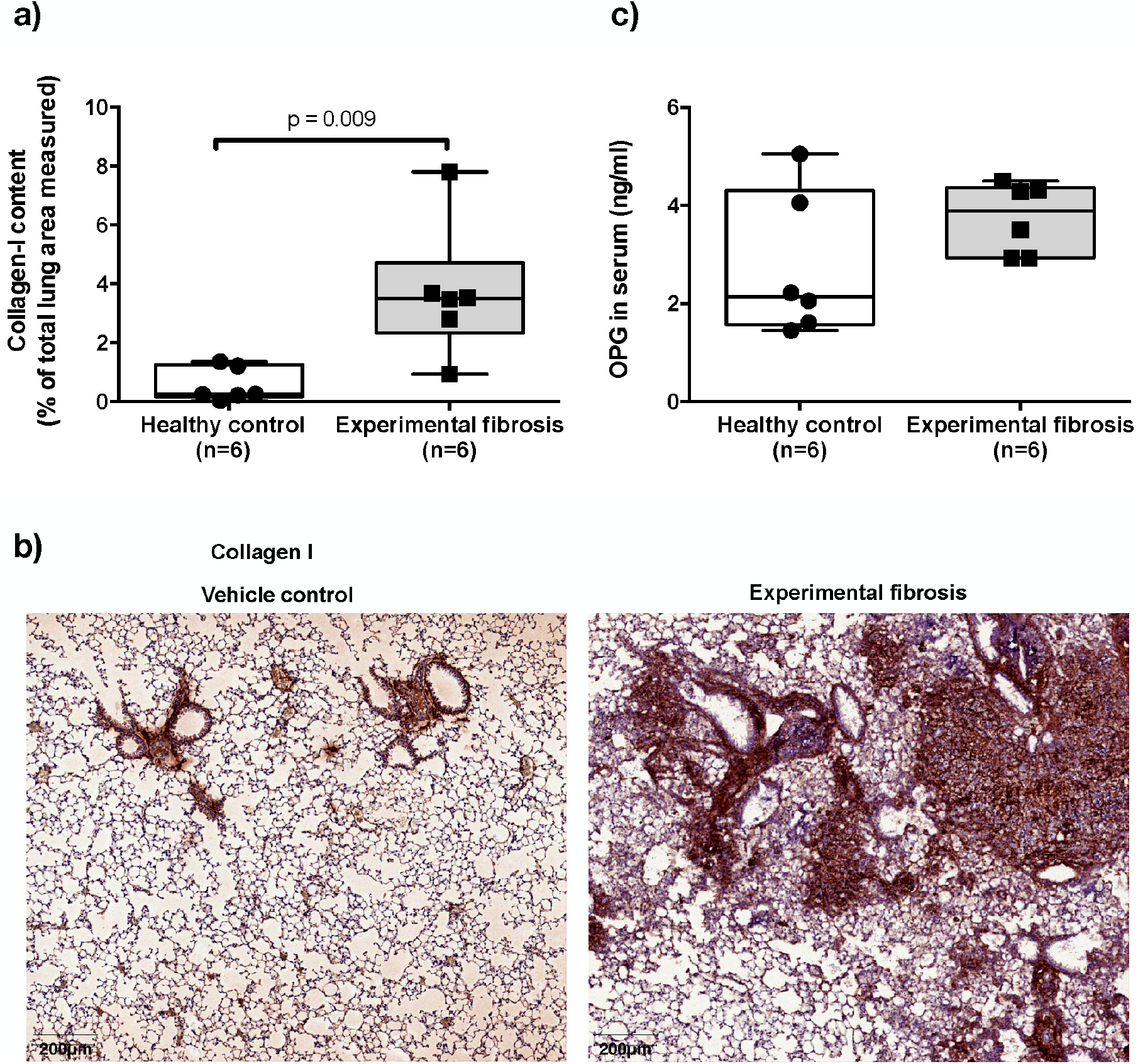
Collagen I is abundantly present in the lung tissue of mice with silica-induced fibrosis. C57BL/6 mice were treated intratracheally with saline or Min-U-Sil 5 crystalline silica and sacrificed after 6 weeks. (a) Collagen I protein content was higher in lungs of mice exposed to silica (fibrosis, n=6) as compared to healthy controls (n=6). (b) Representative pictures of collagen I deposition, showing that collagen I deposition was higher in the lungs of mice exposed to silica as compared to saline-exposed controls. Collagen I expression is indicated by the red color. (c) OPG levels in serum were not significantly different between control (n=6) and silica-exposed mice (n=6). Differences between groups were tested with a Mann Whitney U test. p<0.05 was considered statistically significant.

**Supplemental data figure 2.**
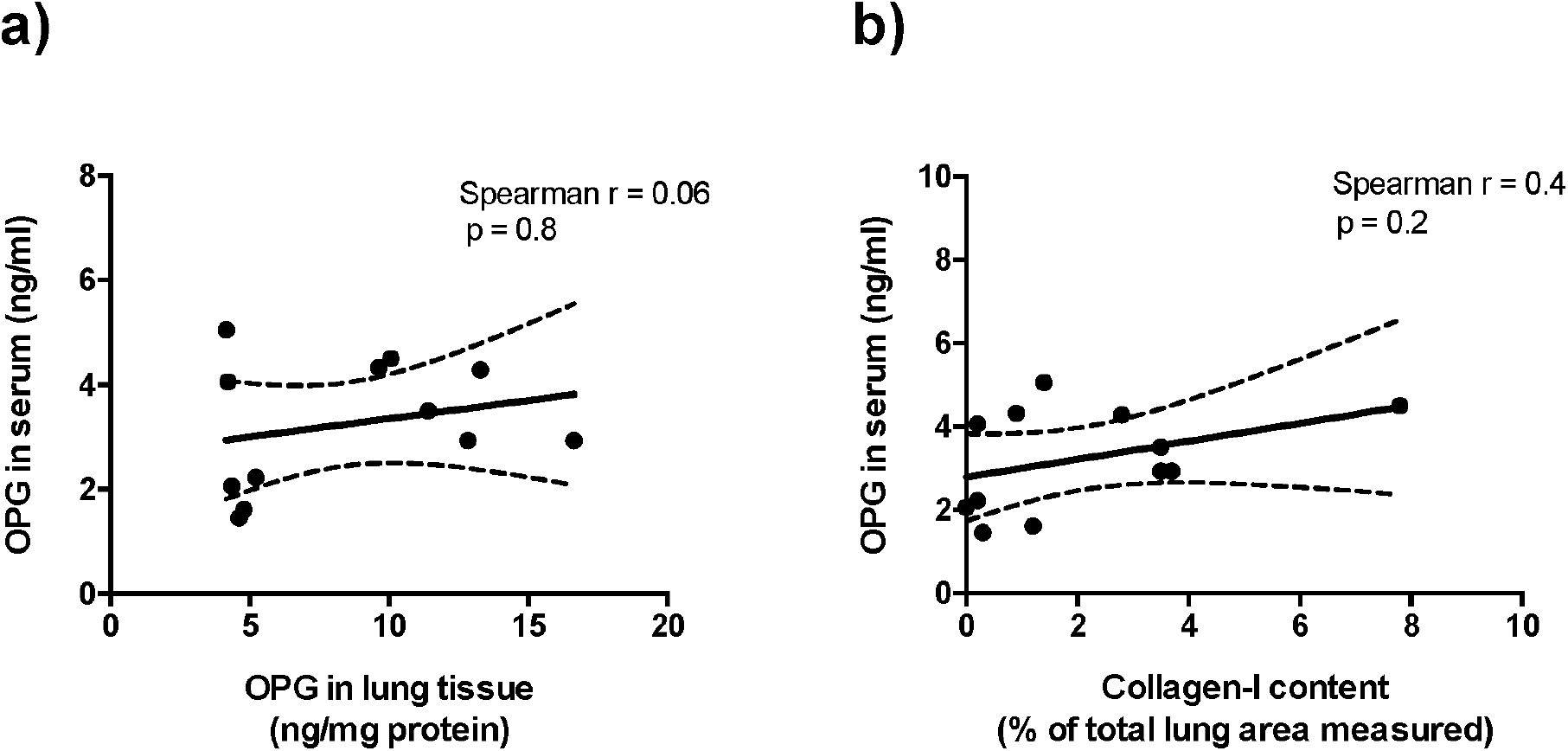
OPG levels in serum do not correlate with OPG levels or collagen I content in lung tissue of control mice and mice with silica-induced fibrosis. C57BL/6 mice were treated intratracheally with saline (n=6) or Min-U-Sil 5 crystalline silica (n=6) and sacrificed after 6 weeks. OPG levels in serum/tissue and collagen I content were determined using ELISA and immunohistochemistry respectively. (a) OPG serum levels do not correlate with OPG levels in mouse lung tissue. (b) OPG serum levels do not correlate with collagen content in mouse lung tissue. Correlation was calculated using a Spearman test and p<0.05 was considered significant.

**Supplemental data figure 3.**
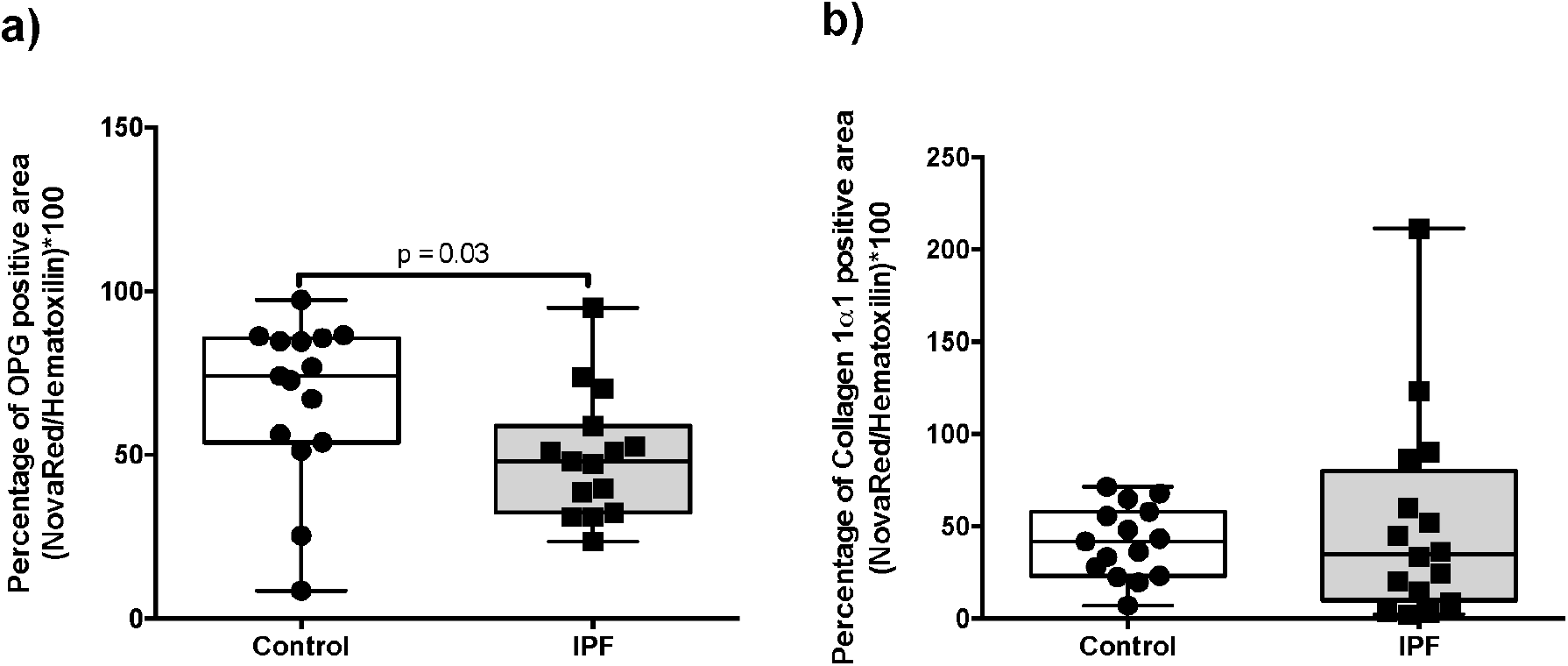
Percentage of OPG and collagen 1α1 positive area in the lung of patients with and without IPF. Lung section of control patients and patients with IPF were stained for OPG and collagen 1α1. Percentage of OPG- (a) and collagen 1α1- positive area (b) in human control and IPF lung tissue was lower in IPF compared to control lung tissue. Differences between groups were tested with unpaired student t test (a) or a Mann Whitney U test (b). p<0.05 was considered statistically significant.

**Supplemental data figure 4.**
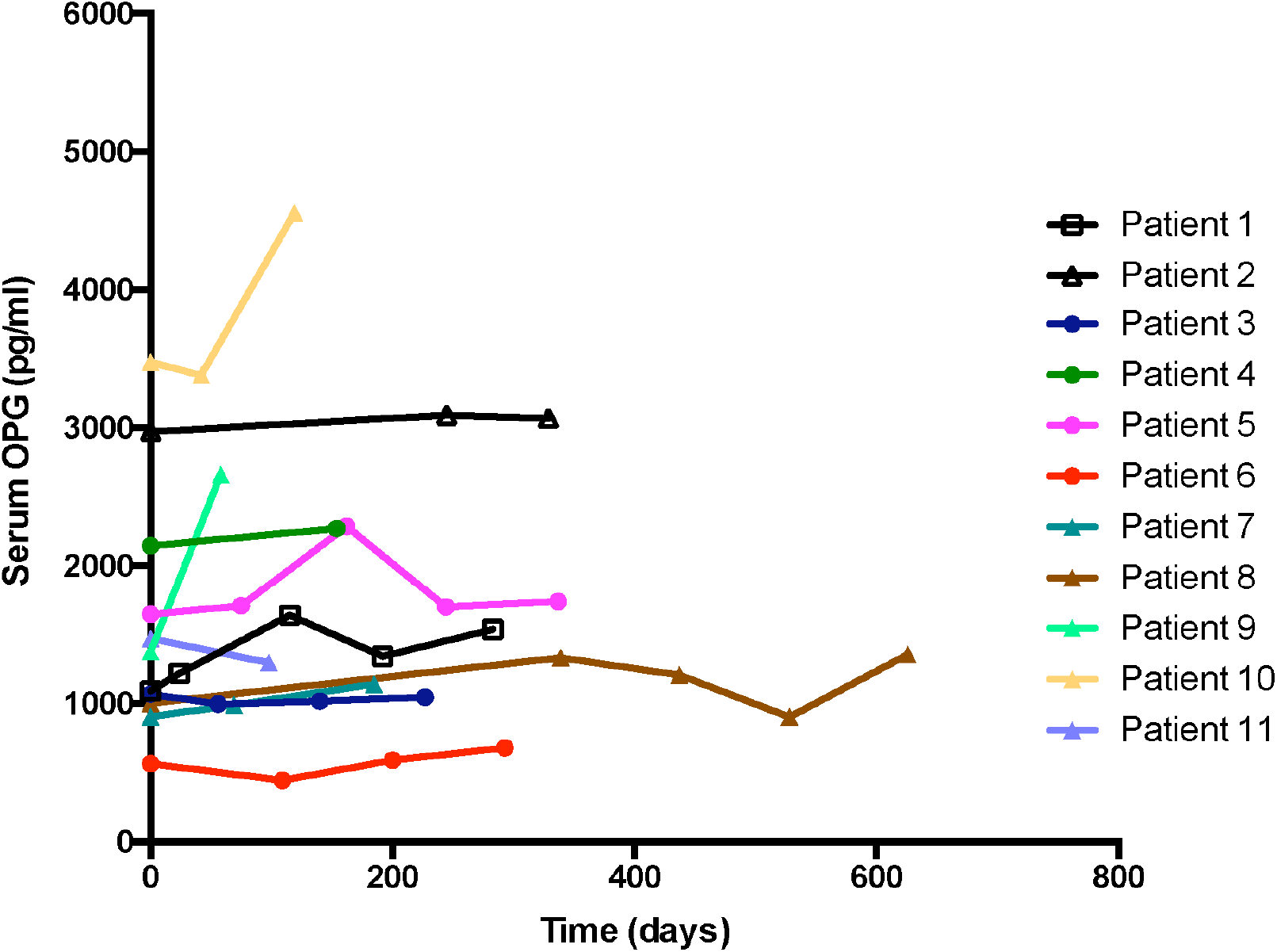
OPG levels did not change over time within patients with IPF. Serum of patients with IPF was collected at multiple time points and OPG levels were measured by ELISA.

